# 3D-printed microfluidic chip for modeling retinal organoid–endothelial co-culture

**DOI:** 10.1101/2025.09.26.678910

**Authors:** Rodi Kado Abdalkader, Shigeru Kawakami, Yuuki Takashima, Takuya Fujita

**Author notes:** Earlier known as Rodi Abdalkader.

## Abstract

Pathological angiogenesis, such as that observed in wet age-related macular degeneration (AMD), is challenging to reproduce in vitro. While previous organ-on-chip approaches have incorporated retinal pigment epithelium (RPE) and endothelial barriers, models integrating human retinal organoids with vascular networks remain limited. Here, we report the development of a fully 3D-printed microfluidic device for co-culture of human induced pluripotent stem cell (hiPSC)-derived retinal organoids containing RPE regions with endothelial cells. The device, fabricated from flexible thermoplastic polyurethane (TPU) on a transparent polyvinyl chloride (PVC) substrate, enables direct organoid–endothelial interaction within a fibrin–Matrigel matrix without physical barriers. In this system, endothelial cells formed choroid-like networks that integrated with retinal organoids. Vascular network density and invasion into RPE regions were enhanced by VEGF stimulation, recapitulating features of wet AMD. Furthermore, fluorescent liposomes distributed along endothelial structures and accumulated at the organoid interface, supporting the application of this model for nanoparticle delivery studies. This 3D-printed retinal organoid-on-chip provides a simple, reproducible, and physiologically relevant platform that complements existing retinal models for investigating angiogenesis and evaluating therapeutic strategies.

## Introduction

Among all eye diseases, retinal diseases can cause severe and irreversible vision loss [1]. Age-related macular degeneration (AMD), diabetic retinopathy, and retinitis pigmentosa are some of the very common illnesses with widespread loss of vision and even blindness[2,3]. The most severe form of AMD, also known as wet AMD, includes abnormal growth of choroidal blood vessels into the subretinal space, causing blurring of vision. The intricate architecture of retinal tissue presents severe challenges to unravelling the pathophysiology of AMD disease aetiology, which in turn hinders effective therapy design.

To advance our knowledge of AMD disease mechanisms, the development of advanced more human-like in vitro models of the retinal tissue is critical. Traditional in vivo models, while valuable, tend to fall short of simulating human retinal pathology due to serious anatomical and genetic differences[4]. Standard in vitro models, on the other hand, can offer controlled environments for the investigation of disease onset, cell interactions, and drug efficacy. However, current in vitro models tend to be less complex and functional compared to the human retinal tissue and thus necessitate more innovative methods for better modeling of retinal disease[5]. Another restriction to the production of in vitro retinal models is that the lack of biological sources for generating retinal cells that closely replicate retinal phenotypes since human donor-derived cells are difficult to acquire, limited to few passages and are costly.

To overcome these issues, we shifted our attention to two major platforms. First, human induced pluripotent stem cell (hiPSC)-based retinal organoids that have shown great promise since they differentiate into various cell types and into retinal structure with a topology simulating native retina—an outstanding progress in retinal modelling[6]. However, these organoids typically lack endothelial cells, which are critical to vascularization processes involved in delivery of nutrients, waste product removal, and recapitulation of pathologies of angiogenesis in disease conditions like AMD. And secondly, the greatest advance in retinal modeling is organ-on-chip (OoC) technology [7,8], which is fast emerging as a strong alternative for establishing more physiologically relevant in vitro models of various organs like the liver [9], the intestine [10], brain [11], and the human eye [12–14]. OoC technology employs microfluidic devices to replicate the microarchitecture and physiology of human organs by placing living cells within a three-dimensional (3D) environment and dynamic flow conditions. In the context of retinal disease research, OoC has several advantages, including its ability to model the blood-retina barrier, provide mechanical and biochemical stimuli by the incorporation of extracellular matrix (ECM), and ability to observe cellular response in real-time manner. These systems can imitate closely the in vivo setting and are hence a better model for studying retinal disease as well as the screening of new therapeutic modalities. Retinal models in OoC were first constructed in a simple setting by emphasizing the importance of the retinal pigment epithelium (RPE) and endothelial cells within the microfluidic devices[15]. The co-culture of hiPSC-derived retinal organoids and RPE cells within the microfluidic devices where direct interaction between these cells was reported as well[16]. While these studies demonstrate the potential of OoC in vision research, direct co-culture of hiPSC-derived retinal organoids with endothelial networks remains limited. Most existing systems use cell lines or rely on layered constructs rather than enabling spontaneous vascular–organoid interactions.

We have previously demonstrated and created a simple microfluidic device via rapid prototyping using fused deposition modeling (FDM) 3D printing of thermoplastic polyurethane (TPU) polymer on a polyvinyl chloride (PVC) substrate[17]. The resulting devices were biocompatible with both endothelial cells and hiPSC-derived optic vesicle organoids, precursors of retinal organoids. This method offers advantages over conventional fabrication techniques, including flexibility, optical transparency, reduced cost, and faster prototyping. In this study, we build on that approach by developing a retinal OoC platform that enables direct co-culture of hiPSC-derived retinal organoids containing RPE regions with endothelial cells within a fibrin–Matrigel matrix. This fully 3D-printed device eliminates the need for semipermeable membranes and allows organoid–endothelial interactions in a physiologically relevant configuration. We show that endothelial cells form choroid-like vascular networks that associate with retinal organoids, and that vascular invasion is enhanced by VEGF stimulation, mimicking features of wet AMD. We further demonstrate that fluorescent liposomes distribute along endothelial networks and localize to the organoid interface, highlighting the suitability of the platform for nanoparticle delivery studies. Together, these results establish a simple, reproducible, and accessible fabrication strategy that complements existing retinal-on-chip systems and broadens their application to studies of angiogenesis and therapeutic testing.

## Results

### 3D-printed microfluidic device provides a flexible and transparent platform for retinal organoid culture

To develop a physiologically relevant co-culture platform for retinal organoids and endothelial cells without the need for a physical barrier such as a porous membrane (**Figure 1**), we fabricated a multi-well microfluidic device from flexible TPU using FDM 3D printing on a transparent PVC substrate, enabling optical monitoring. Each unit contained six culture chambers, with a central compartment for organoid culture and two lateral reservoirs for media exchange. A schematic cross-section illustrates the chamber dimensions and media flow channels (**Figure 2a**). Perfusion tests with fluorescein dye confirmed leak-free connectivity across the device (**Figure 2b**). SEM images revealed well-defined filament deposition, consistent interlayer adhesion, and intact chamber geometry. The transparency of the PVC supported high-resolution imaging, while the flexibility of the TPU facilitated repeated handling during culture (**Figure 2c**; **Supplementary Figure 1**).

**Figure 1.**
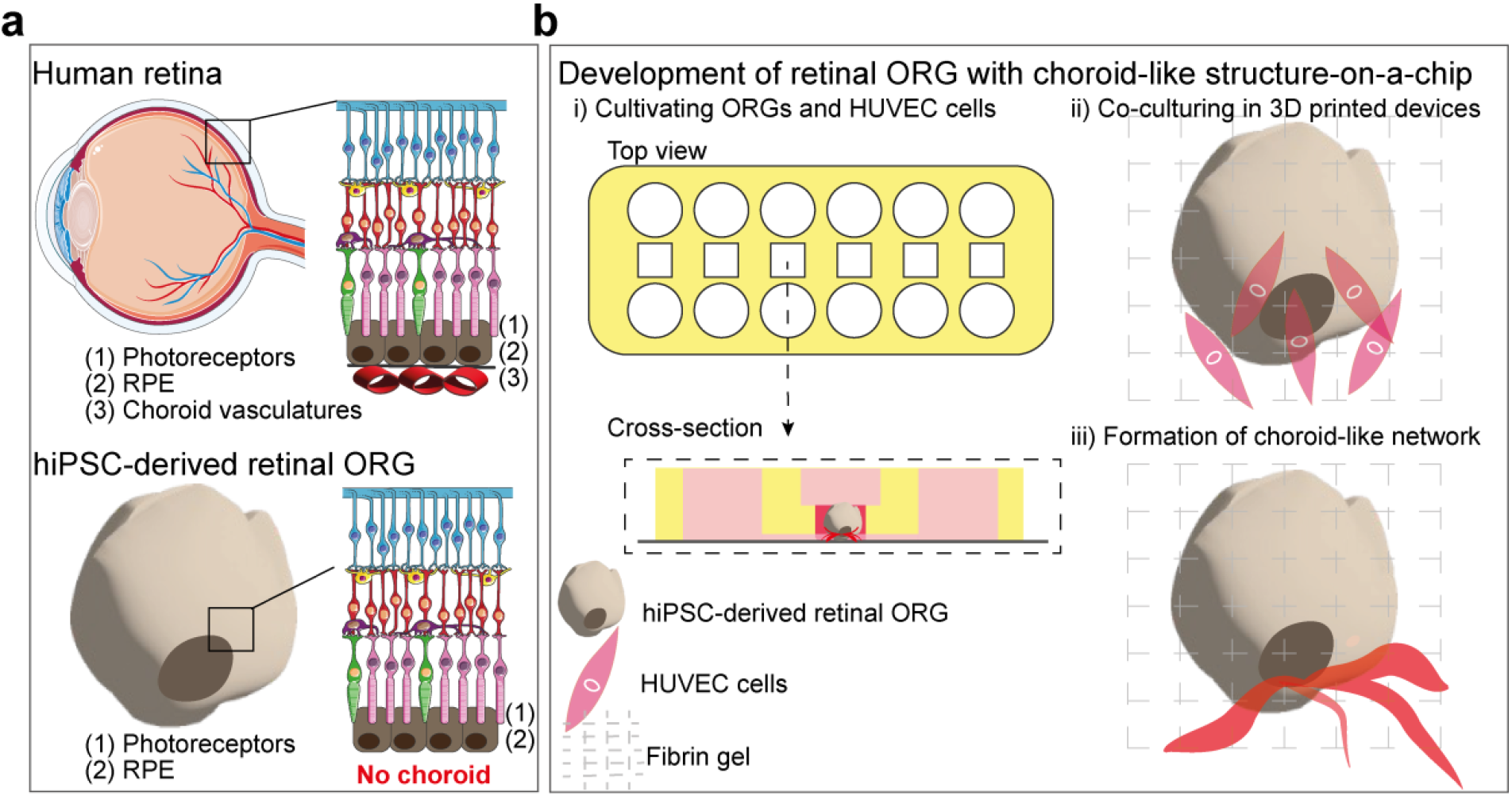
Schematic representation of the human retina and the development of a retinal organoid with a choroid-like structure-on-a-chip. (a) Comparison of the native human retinal structure (top) with hiPSC-derived retinal organoids (ORG) (bottom). The human retina consists of photoreceptors (1), retinal pigment epithelium (RPE; 2), and underlying choroidal vasculature (3). In contrast, hiPSC-derived retinal organoids recapitulate photoreceptors (1) and RPE (2) but lack choroidal vasculature. (b) Strategy for developing a retinal organoid with a choroid-like structure in a 3D-printed microfluidic device. (i) Retinal organoids and HUVECs are cultivated and introduced into the device. Top and cross-sectional views show organoids embedded in fibrin gel adjacent to endothelial cells. (ii) Endothelial cells are co-cultured with retinal organoids, enabling interaction at the RPE border. (iii) Endothelial cells form a choroid-like vascular network surrounding the organoid, mimicking native retinal–choroidal interactions. Part of the illustration of the retina and eye was adapted from Servier Medical Art (https://smart.servier.com/), licensed under CC BY 4.0 (https://creativecommons.org/licenses/by/4.0/).

**Figure 2.**
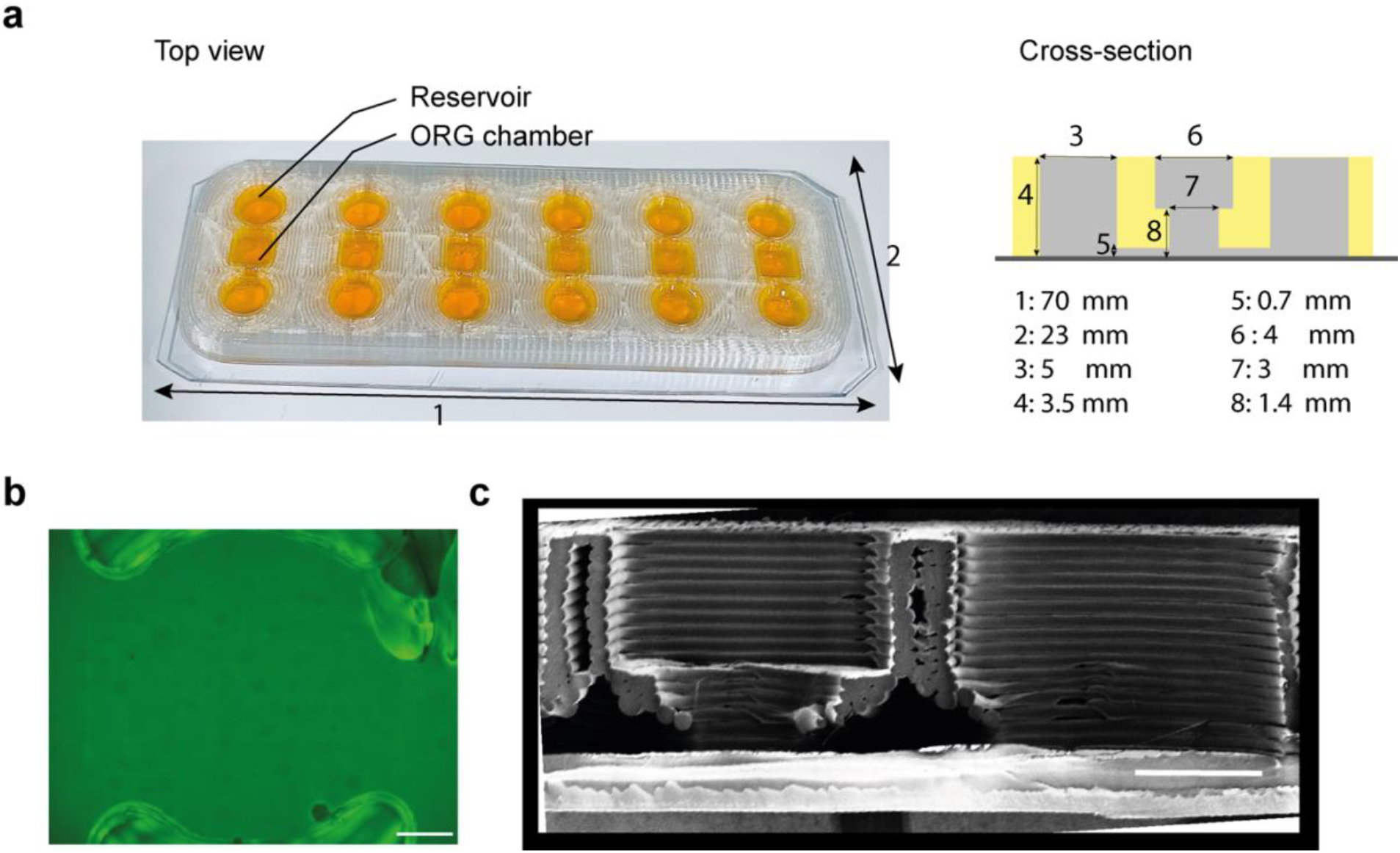
Design and characterization of the 3D-printed microfluidic device. (a) Top view of the fabricated device showing reservoirs and organoid-endothelial cells placing chambers (left), and schematic cross-section with corresponding dimensions (right). (b) Fluorescence image showing flow of fluorescein solution in the organoid chamber, demonstrating liquid connectivity across the device. Scale bar: 500 µm. (c) SEM cross-sectional image of the printed device, revealing the layered structure of the TPU filaments and embedded chamber geometry. Scale bar: 1000 µm.

### Endothelial cells self-organize into vascular networks around retinal organoids

To develop choroid-like vasculature, hiPSC-derived retinal organoids with RPE were co-cultured with human umbilical vein endothelial cells (HUVECs) in a 3D-printed microfluidic device. Organoids were generated over 93 days, followed by a 5-day co-culture period within the microfluidic platform (**Figure 3a**). In this system, HUVECs were embedded in fibrin gel adjacent to the organoid, promoting the formation of vascular networks. 3D reconstruction of the vascular networks revealed significant differences between monoculture and co-culture conditions (**Figure 3b**). In monoculture, HUVECs formed sparse and thin networks. In contrast, the co-culture condition with organoids (H_ORG) resulted in a wider interconnected and organized network. Quantitative analysis of the mean fluorescence intensity (MFI) across the z-axis showed that the monoculture networks were primarily localized at the bottom layer, whereas in the co-culture condition, MFI was distributed more uniformly throughout the construct (**Figure 3c**). This suggests enhanced vascular integration and penetration in the presence of organoids. Confocal imaging confirmed the structural organization and integration of the vascular networks with organoids (**Figure 3d**). The networks exhibited extensive connections with the RPE regions in organoids, demonstrating spatial integration and potential physiological relevance of the choroid-like networks.

**Figure 3.**
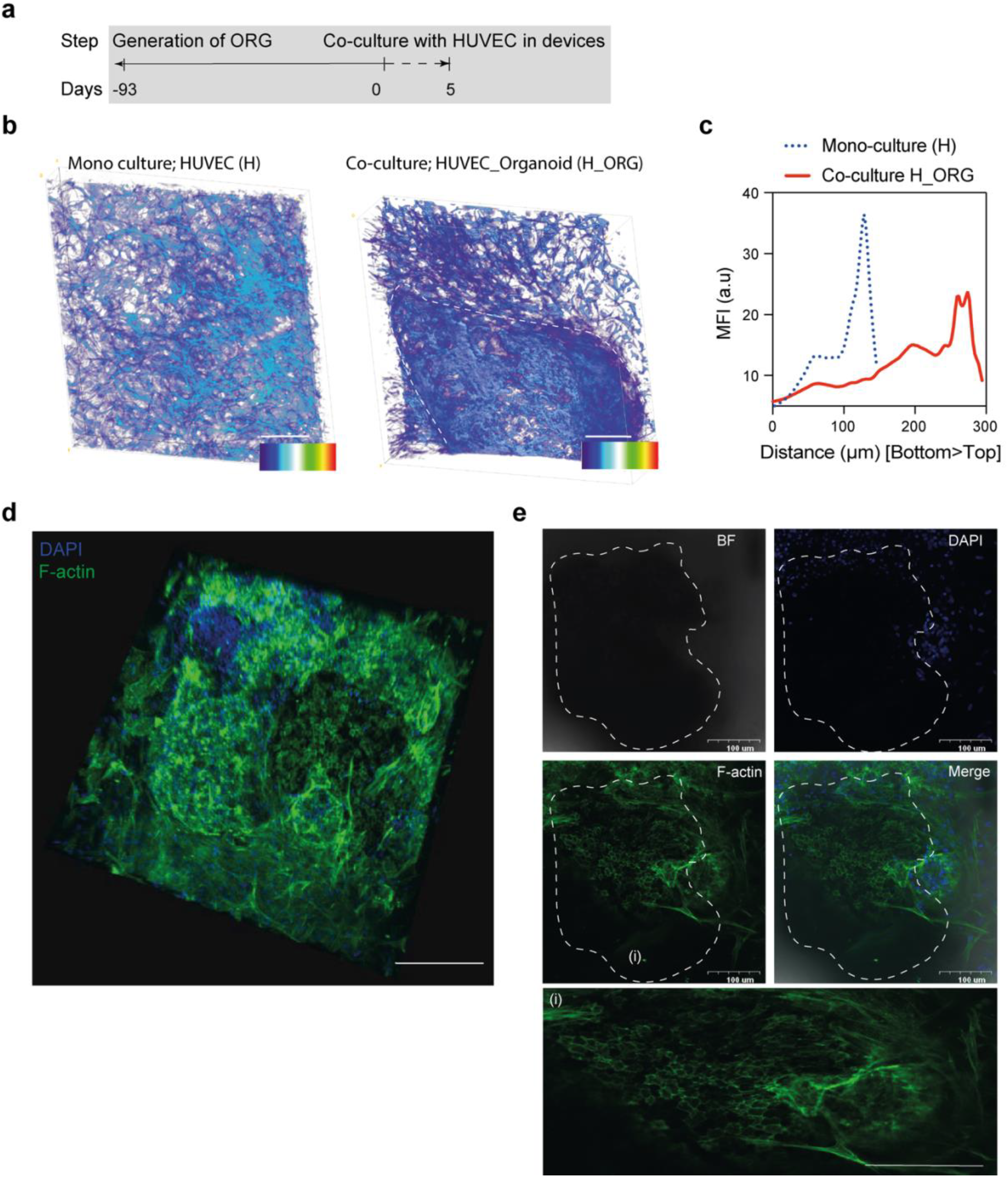
Vascular network formation of HUVECs in co-culture with retinal organoids. (a) Timeline of experimental design showing retinal organoid (ORG) generation and subsequent co-culture with HUVECs in microfluidic devices. (b) Representative 3D reconstructed images of HUVEC monoculture (H) and co-culture with retinal organoids (H_ORG), showing distinct endothelial organization. (c) Mean fluorescence intensity (MFI) profiles across z-stacks, comparing monoculture (dotted blue) and co-culture (red) conditions. (d) Confocal image of endothelial networks in co-culture stained for F-actin (green) and nuclei (DAPI, blue), demonstrating extensive vascular-like structures. Scale bar: 200 µm. (e) Immunofluorescence analysis of organoid–endothelial co-culture. Bright-field (BFI), DAPI, F-actin, and merged images (top panels) highlight endothelial localization around the retinal organoid (outlined by dashed lines). Magnified view (i) shows endothelial network extension at the RPE-organoid interface. Scale bars: 100 µm (top), 100 µm (i).

### VEGF stimulation enhances endothelial invasion and preserves retinal organoid identity

To model pathological angiogenesis, VEGF (300 ng mL^−1^) was added during the 7-day co-culture of retinal organoids and HUVECs within the 3D-printed device (**Figure 4a**). To evaluate the connectivity of endothelial structures, FITC-labeled beads were introduced into the chamber. Bright-field and fluorescence imaging showed the distribution of beads along endothelial networks (**Figure 4b**). Quantitative analysis of MFI from the center to the periphery of the chamber revealed a gradual increase in signal toward the edges (**Figure 4c**), consistent with directed bead distribution within the vascular-like structures. Confocal imaging further confirmed the organization of endothelial cells in proximity to the organoid. Networks stained for F-actin formed dense structures that extended toward and contacted organoid borders, with clear association at RPE-rich regions (**Figure 4d**). This demonstrated that VEGF stimulation promoted endothelial sprouting and invasion into organoid niches, resembling features of wet AMD. Histological analysis further supported these findings. Hematoxylin and eosin (H&E) staining of paraffin sections revealed two distinct regions: the retinal organoid (i) and the surrounding endothelial-rich area (ii), with visible interactions at their interface (**Figure 4e**). This structural evidence reinforced the confocal observations of endothelial integration into the organoid environment. To verify that retinal identity was preserved during endothelial co-culture, immunofluorescence staining was performed for key retinal markers. Organoids expressed the neuronal marker TUJ1 and the retinal marker VSX2, indicating that they maintained layered neuronal and retinal cell populations under co-culturing conditions (**Figure 4f**).

**Figure 4.**
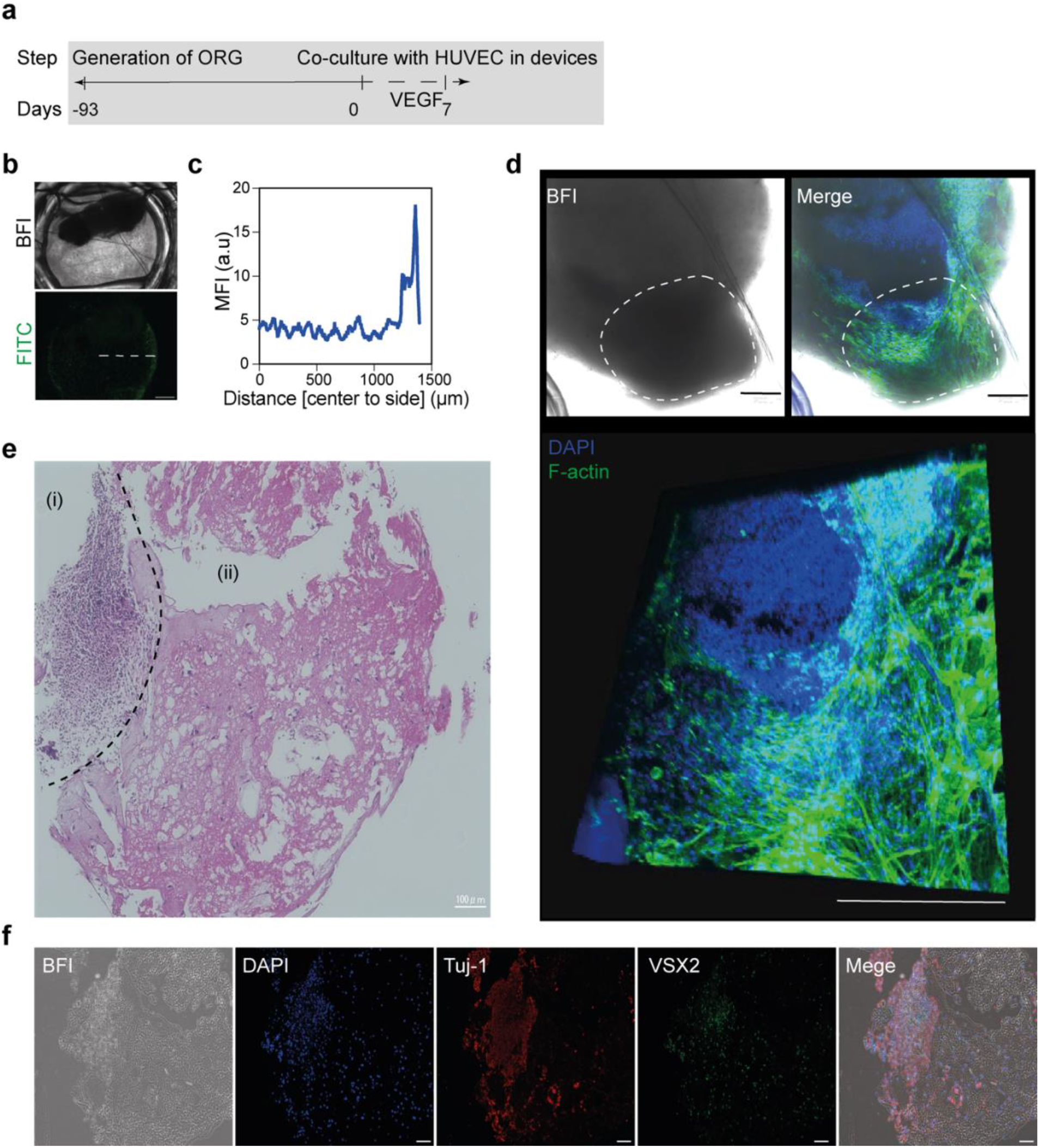
VEGF-induced angiogenesis and structural characterization of retinal organoid–endothelial co-culture. (a) Timeline of experimental design showing retinal organoid (ORG) generation, co-culture with HUVECs in devices, and VEGF treatment for 7 days. (b) Bright-field (BFI) and fluorescence (FITC) images showing FITC-labeled bead flow through the co-culture chamber, demonstrating functional perfusability of the microfluidic channels. (c) Mean fluorescence intensity (MFI) along the chamber axis (center-to-side profile), confirming bead distribution and flow. (d) Confocal images of endothelial networks in co-culture stained with F-actin (green) and nuclei (DAPI, blue). BFI and merged views, with dashed lines outlining the retinal organoid boundary. Scale bar: 200 µm. (e) Hematoxylin and eosin (H&E) staining of organoid– endothelial co-culture section. (i) Retinal organoid region and (ii) surrounding endothelial-rich area. Scale bar: 100 µm. (f) Immunofluorescence analysis of retinal organoid differentiation within the co-culture. Representative images of BFI, nuclei (DAPI), neuronal marker Tuj-1 (red), retinal progenitor marker VSX2 (green), and merged overlay demonstrate preservation of retinal identity. Scale bar: 100 µm.

### Nanoparticle liposomes distribute through endothelial networks and accumulate at the organoid interface

To test whether endothelial networks could mediate nanoparticle transport, rhodamine-labeled liposomes were introduced into the inlet chamber (**Figure 5a**; **Supplementary table 2**). Time-lapse imaging showed progressive accumulation of rhodamine signal near organoids at 0, 10, 20, and 30 min (**Figure 5b**). Quantitative MFI profiles confirmed a time-dependent increase in fluorescence close to the organoid, with maximal enrichment at 30 min (**Figure 5c**).

**Figure 5.**
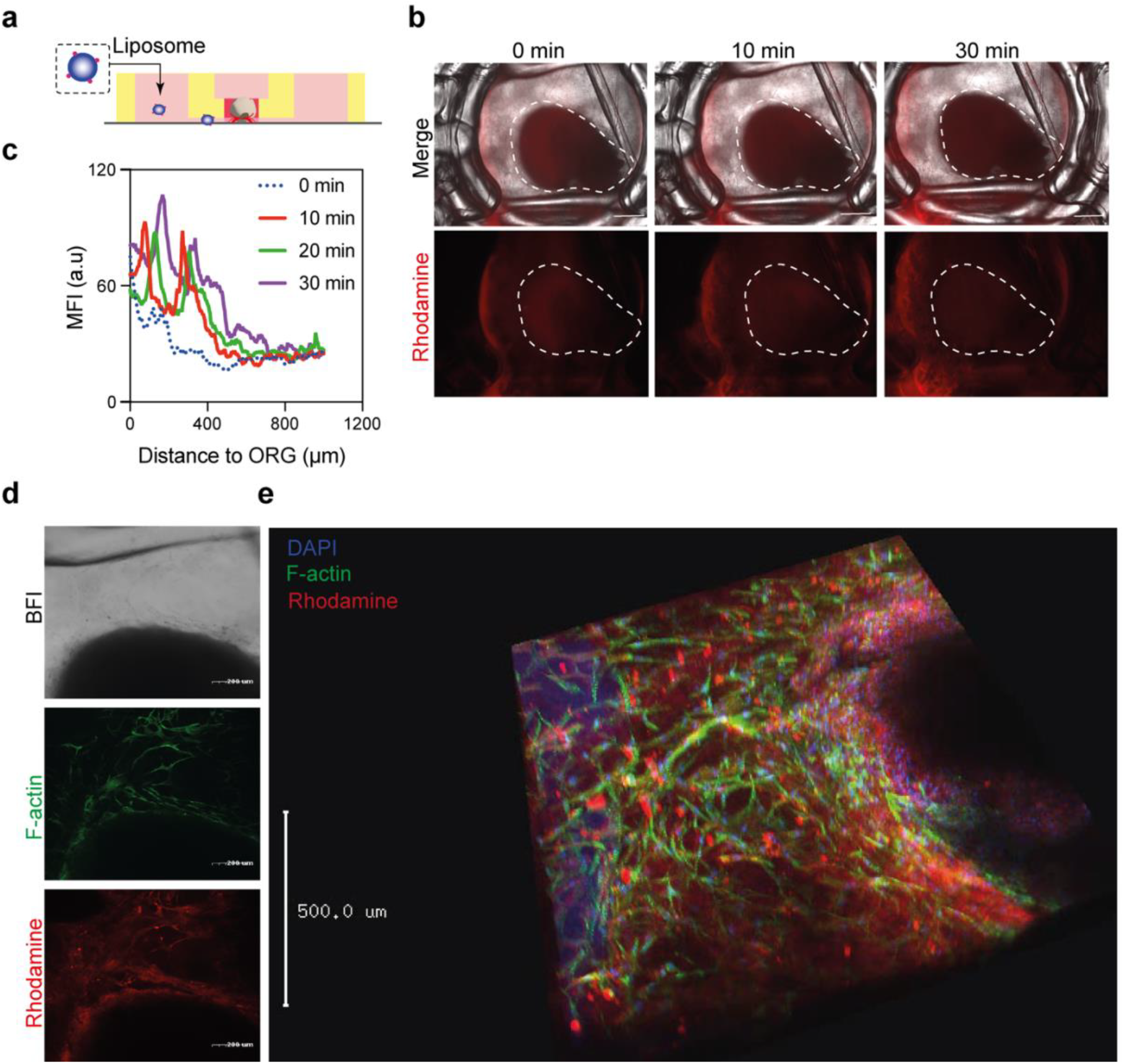
Liposome delivery in retinal organoid–endothelial co-culture. (a) Schematic illustration of liposome delivery through the microfluidic device into the retinal organoid chamber. (b) Time-lapse bright-field and fluorescence images showing rhodamine-labeled liposome perfusion at 0, 10, 20, and 30 min. Dashed lines outline the retinal organoid. (c) Mean fluorescence intensity (MFI) profiles at 0, 10, 20, and 30 min along the chamber axis, demonstrating progressive accumulation of rhodamine signal near the organoid (ORG). (d) BFI, F-actin (green), and rhodamine (red) images showing liposome uptake and endothelial association in the co-culture. Scale bar: 200 µm. (e) 3D confocal reconstruction of co-cultures stained with F-actin (green), DAPI (blue), and rhodamine-labeled liposomes (red), confirming vascular network perfusion and delivery to the organoid microenvironment. Scale bar: 500 µm.

To visualize nanoparticle localization relative to endothelial cells and organoid tissue, confocal imaging was performed after 60 min of liposome exposure. Bright-field and fluorescence images revealed endothelial networks (F-actin, green) adjacent to organoids, with rhodamine-positive liposomes (red) colocalized along endothelial structures and at the organoid boundary (**Figure 5d**). A 3D confocal reconstruction confirmed distribution of liposomes throughout the vascular-like network and accumulation at the organoid interface (**Figure 5e**). Together, these findings demonstrate that the engineered endothelial networks not only form choroid-like structures and invade organoid RPE regions under VEGF stimulation but also mediate distribution of nanoparticle carriers toward retinal tissue.

## Discussion

The integration of endothelial cells with retinal organoids is essential for replicating the abnormal vascularization characteristic of wet AMD. In this study, we developed an organ on chip (OoC) system using a 3D-printed device that enables the co-culture of endothelial cells with hiPSC-derived retinal organoids containing RPE regions embedded in fibrin gel.

We employed our previously established method for rapid and straightforward device prototyping using FDM 3D printing. Moreover, in our earlier studies, we demonstrated that the use of flexible TPU and PVC materials exhibits high compatibility with human iPSC-derived optical vesicles, allowing them to maintain and preserve their characteristics. This approach was adopted for the fabrication of the OoC system, offering several advantages: seamless one-step fabrication, the ability to incorporate ECM gel materials, and an array of six sample wells, enabling multiple parallel experiments.

To investigate the formation of choroid-like networks of HUVEC cells with ORG-RPE within fibrin gel, we employed a basal medium free from VEGF. Our results indicated that the co-culture has improved the vasculature networking as compared to monoculture presumably due to the impact of secreted factor from RPE cells. Moreover, the addition of VEGF into the culturing medium has significantly accelerated the formation of dense vasculature network that totally covered and invaded the RPE spots in organoids, a characteristic feature of wet AMD.

In similar applications, previous studies have reported the development of retinal barrier models using RPE cell lines and endothelial cells. However, these models primarily relied on cell lines such as RPE cells [18], or involved placing organoids on top of an RPE monolayer rather than integrating them as a complete system [16]. In our study, we utilized hiPSC-derived retinal organoids with intrinsic RPE regions and co-cultured them with endothelial cells to investigate the angiogenesis process. This was examined under both basal conditions and in response to high concentrations of VEGF

Despite these advantages, several limitations should be acknowledged. In our 3D culture system, the extracellular matrix acts as a barrier between endothelial and RPE cells, while also permitting endothelial interactions with photoreceptors in regions not covered by RPE. Furthermore, HUVECs may not fully recapitulate the properties of native choroidal endothelial cells. Incorporating hiPSC-derived or primary choroidal endothelial cells in future studies will be critical to enhance model fidelity.

Taken together, our work establishes a foundation for unified co-culture systems in which RPE, endothelial cells, and retinal organoids coexist within a single device. Such systems will enable direct cell–cell communication and more accurately mimic the native retinal microenvironment. Future refinements in endothelial cell sourcing and device architecture could yield highly physiologically relevant and personalized models for studying retinal angiogenesis and for developing therapeutic strategies for wet AMD.

### Conclusion

In this study, we established a fully 3D-printed retinal organoid-on-a-chip platform that enables direct co-culture of hiPSC-derived retinal organoids containing intrinsic RPE regions with endothelial cells. By eliminating semipermeable barriers and embedding cells within a fibrin–Matrigel matrix, this system allows physiologically relevant organoid–endothelial interactions. We demonstrated that endothelial cells self-organized into choroid-like vascular networks that associated with retinal organoids, and that VEGF stimulation promoted sprouting and invasion into RPE regions, recapitulating key features of wet AMD. Furthermore, the device supported nanoparticle liposome transport through engineered vascular networks, highlighting its utility for drug delivery and therapeutic testing.

This simple and reproducible platform offers a promising tool for studying retinal angiogenesis and testing therapeutic strategies.

## Methods

### Device fabrication and characterization

The microfluidic device was fabricated using our previously established FDM 3D-printing protocol (Anycubic Technology Co., Ltd., Shenzhen, China) with 1.75 mm translucent clear TPU filaments (SainSmart, Lenexa, KS, USA)[17,19]. Device designs were generated in Tinkercad (https://www.tinkercad.com) and consisted of six cell culture chambers, each connected to inlets and outlets. STL files were sliced and processed using Ultimaker Cura software with the following parameters: layer height, 0.01–0.02 mm; shell thickness, 2–10 layers. TPU filaments were directly deposited onto 1 mm-thick clear PVC sheets, after which the printed constructs were trimmed and stored at room temperature. Structural characterization of the printed devices was performed by scanning electron microscopy (SEM; VE-9800, Keyence Corp., Osaka, Japan). To evaluate liquid flow within the microfluidic channels, a sodium fluorescein solution (1 mg mL^−1^; Nacalai Tesque Inc., Kyoto, Japan) and fluorescent yellow, green-labeled latex microbeads (2µm; Sigma-Aldrich, St. Louis, MO, USA) were perfused through the inlet–chamber–outlet pathway, allowing visualization of particle transport across the device.

### Cell culturing and organoid generation

hiPSCs, 585A1[20], was procured from Riken Cell Bank and handled in accordance with the guidelines stipulated by the ethics committees of Ritsumeikan University (Approved number 2021-004-3). Culture dishes were pre-coated with imatrix-511 (Nippi Inc., Tokyo, Japan) overnight at 37°C. 1×10^4^ viable cells in mTeSR^TM^ plus medium (STEMCELL Technologies Inc., Vancouver, Canad**a**) supplemented with 10 µM Y27632 (FUJIFILM Wako, Osaka, Japan) were seeded in 6-well plate. The next day cell culture medium was changed to mTeSR^TM^ plus only. The culture medium was refreshed with fresh mTeSR^TM^ plus medium every two days. Subsequently, embryoid bodies (EBs) were produced by adding 1×10^5^ viable cells in mTeSR^TM^ plus medium supplemented with 10 µM Y27632 in 10 cm ultra-low attachment dishes (PrimeSurface®; Sumitomo Co., Ltd., Tokyo, Japan). After three days, the mTeSR^TM^ plus medium was replaced with the chemical induction medium, consisting of fresh E6 medium (Thermo Fisher Scientific Inc., Waltham, MA, USA) supplemented with 10 µM TGF-β kinase/activin receptor-like kinase inhibitor (SB-505124; Santa Cruz Biotechnology Inc., Dallas, TX, USA), 10 µM Y27632 and BMP4 (20 ng mL^-1^)(PeproTech, Rocky Hill, NJ, USA). These conditions were maintained for 3 days. EBs were then transferred into 6-well plates coated with matrigel (Geltrex; Thermo Fisher Scientific Inc., Waltham, MA, USA) (6-8 EBs per well). Subsequently, the medium was switched to E6 medium supplemented with 500 nM retinoic acid (FUJIFILM Wako, Osaka, Japan) for 7 days. On day 20, optical vesicle (OV) organoids aggregates were picked with 27 G syringe needle (Terumo Corp., Tokyo, Japan) and transferred in E6 medium supplemented with 10 µM Y27632 and 100 µM taurine to 10 cm ultra-low attachment dishes (PrimeSurface®; Sumitomo Co., Ltd., Tokyo, Japan). The culture medium was refreshed with E6 medium supplemented 100 µM taurine (Sigma-Aldrich, St. Louis, MO, USA) every two days. From day 50, 500 nM of retinoic acid (FUJIFILM Wako, Osaka, Japan) was added to the medium for inducing maturation.

For culturing HUVEC culturing, cells were passaged on 0.1 % gelatin pre-coated dished in DMEM medium supplemented with 10 % v/v FBS, 10 ng mL^-1^ bFGF (FUJIFILM Wako, Osaka, Japan) and 1ng mL^-1^ EGF (FUJIFILM Wako, Osaka, Japan) to be known as HUVEC basal medium. The medium was replaced, and the culture medium was refreshed daily.

### Cell co-culture and VEGF-induced angiogenesis assay

HUVECs were suspended in an embedding gel consisting of 100 µL bovine fibrinogen (5 mg mL^-1^; FUJIFILM Wako, Osaka, Japan), 320 µL Matrigel (Geltrex; Thermo Fisher Scientific, Waltham, MA, USA), and 4 µL aprotinin (FUJIFILM Wako, Osaka, Japan). Separately, 1 × 10^7^ cells were resuspended in 425 µL DMEM and then mixed with the embedding gel to yield the final cell–gel mixture, which was kept on ice. Microfluidic devices were sterilized by UV irradiation for 60 min before use. Retinal organoids were transferred to the central chamber of each device using a wide-bore pipette tip, overlaid with 40 µL of the cell–gel mixture, and immediately treated with 10 µL thrombin (5 U mL^-1^; FUJIFILM Wako, Osaka, Japan). After incubation at 37 °C with 5% CO_2_ for 10 min to allow gel polymerization, the medium was replaced with HUVEC basal medium supplemented with 1 µg mL^-1^ aprotinin. To induce angiogenesis, VEGF was added to the HUVEC basal medium at a final concentration of 300 ng mL^-1^ (FUJIFILM Wako, Osaka, Japan) and applied daily to the device reservoirs. For the nanoparticle delivery assay, rhodamine-labeled liposomes were prepared and characterized as described in supplementary Method 1. Liposomes were diluted in Hank’s Balanced Salt Solution supplemented with calcium and magnesium (HBSS^+^; Thermo Fisher Scientific) to a final concentration of 0.1 mg mL^−1^ of lipids and then 10 µL introduced into the inlet of the microfluidic device. Liposome distribution was monitored by time-lapse fluorescence microscopy over a 30 min period (KEYENCE, Tokyo, Japan). After 60 min of exposure, co-cultures were fixed with 4% paraformaldehyde at room temperature for 30 min. Samples were then washed with PBS and imaged by confocal laser scanning microscopy (FV3000; Olympus Corp., Tokyo, Japan).

### Microscopic imaging and structure analysis

Samples were washed with PBS twice and then fixed with 4% paraformaldehyde in PBS-for 60 min at 25°C. Samples were then permeabilized with 0.1% Triton X-100 at 4 ºC for overnight. Subsequently, cells were incubated with ActinGreen 488 (Thermo Fisher Scientific Inc., Waltham, MA, USA) at 4 ºC for overnight. And then scaffolds were washed with PBS twice and then incubated with Hoechst 33342 (Dojindo Molecular Technologies, Inc., Kumamoto, Japan) in a final concentration of 10 µg mL^-1^ for 60 min at 25°C. For imaging, we used confocal scanning microscope (FV3000; Olympus Corp., Tokyo, Japan). Images were then analysed using FV31S-SW software. For the investigation of cell morphology and fluorescence intensity in ImageJ software was employed (National Institute of Health, Maryland, USA).

### Paraffin blocks, H&E and immunofluorescence staining

Organoids were fixed with 4% paraformaldehyde in PBS overnight at 4°C and then processed to generate paraffin blocks. Subsequently, 5 µm slices were obtained and stored at 4°C. For H&E staining, samples were deparaffinized prior to staining. For immunofluorescence staining, samples underwent antigen retrieval using Tris-EDTA pH 9.0, until boiling for 15 min, followed by permeabilization with 0.5% Triton X-100 in PBS overnight at 4°C. The cells were then blocked with blocking buffer (5% bovine serum albumin, 0.1% Tween-20) at room temperature for 90 min and subsequently incubated overnight at 4°C with primary antibodies diluted in blocking buffer (Tuj-1, VSX2 or CRX). After washing, cells were incubated at 37°C for 60 min with the appropriate secondary antibody (Alexa Fluor 555 Goat anti-mouse IgG, Alexa Fluor 488 Goat anti-rabbit IgG; 1:500 v/v; Thermo Fisher Scientific) (**Supplementary table 1**). Finally, cells were mounted with antifade solution supplemented with the nucleus stain DAPI Fluoro-KEEPER (Nacalai, Kyoto, Japan) at 25°C for 60 min. Imaging was performed using a fluorescence microscope (KEYENCE, Tokyo, Japan) to acquire cell images from cell chambers.

## Data visualization and statistics

For the investigation of cell morphology and quantification of Mean fluorescence intensity (MFI), images were analyzed using ImageJ software (National Institutes of Health, Bethesda, MD, USA). Graphs generated using GraphPad Prism 9 (GraphPad Software, San Diego, CA, USA). Data are represented as mean ± S.E.M.

## Acknowledgments

We acknowledge the Ritsumeikan Global Innovation Research Organization (R-GIRO).

## Authors contribution statement

Rodi Kado Abdalkader: Conceptualized and managed the project, designed, and performed experiments, analyzed, and interpreted data, visualized the data, and wrote the manuscript. Shigeru Kawakami, Yuuki Takashima, and Takuya Fujita: provided resources and contributed to manuscript refinement. All authors critically reviewed the manuscript and agreed with the publication.

## Additional information

Competing interests: All authors declare no competing financial interest.

## Data availability

The datasets used and/or analysed during the current study are available from the corresponding author on reasonable request.

