## Supplementary Information for "3D-printed microfluidic chip for modeling retinal organoid–endothelial co-culture"

*Corresponding author.

†Earlier known as Rodi Abdalkader

### Supplementary Methods

#### Preparation of rhodamine-labeled liposomes

Liposomes were prepared as previously described with minor modifications. A total lipid mass of 20 mg was used, consisting of distearoyl-sn-glycero-3-phosphocholine (DSPC; Avanti Polar Lipids, Alabaster, AL, USA), 1,2-distearoyl-sn-glycero-3-phosphoethanolamine-N-[amino(polyethylene glycol)-2000] (DSPE-PEG2000; NOF Corporation, Tokyo, Japan), and 1,2-dioleoyl-sn-glycero-3-phosphoethanolamine-N-(lissamine rhodamine B sulfonyl) (Rhodamine-DHPE; Thermo Fisher Scientific, Waltham, MA, USA) in a 93:6:1 molar ratio. Lipids were dissolved in chloroform, and the solvent was evaporated at 25 °C, followed by vacuum drying overnight at room temperature. The resulting lipid film was hydrated with phosphate-buffered saline (PBS, pH 7.4) pre-warmed to 65 °C for 60 min under mild agitation, giving a final lipid concentration of 8 mg $mL^{-1}$. The suspension was sonicated in a bath sonicator for 10 min, followed by probe sonication for 3 min, yielding small unilamellar vesicles. Liposomes were stored at 4 °C protected from light until use. Dynamic light scattering (DLS; Zetasizer Nano ZS, Malvern Instruments, UK) was used to measure average hydrodynamic diameter.

**Supplementary table 1.**

| Target protein | Host species | Maker | Catalog # | Working dilution |
|---|---|---|---|---|
| Beta Tubulin 3/ Tuj1 | Mouse | Gene Tex | GTX27751 | 1:250~ |
| CRX | Rabbit | Proteintech | 12047-I-AP | 1:100~ |
| VSX2 | Rabbit | Proteintech | 25825-1-AP | 1:100~ |

**Supplementary table 2.**

| DSPC: $DSPEPEG_{2000}$: Rhodamine-DHPE | |
|---|---|
| Size | PDI |
| 77.69 ± 2.17 | 0.27 ± 0.081 |

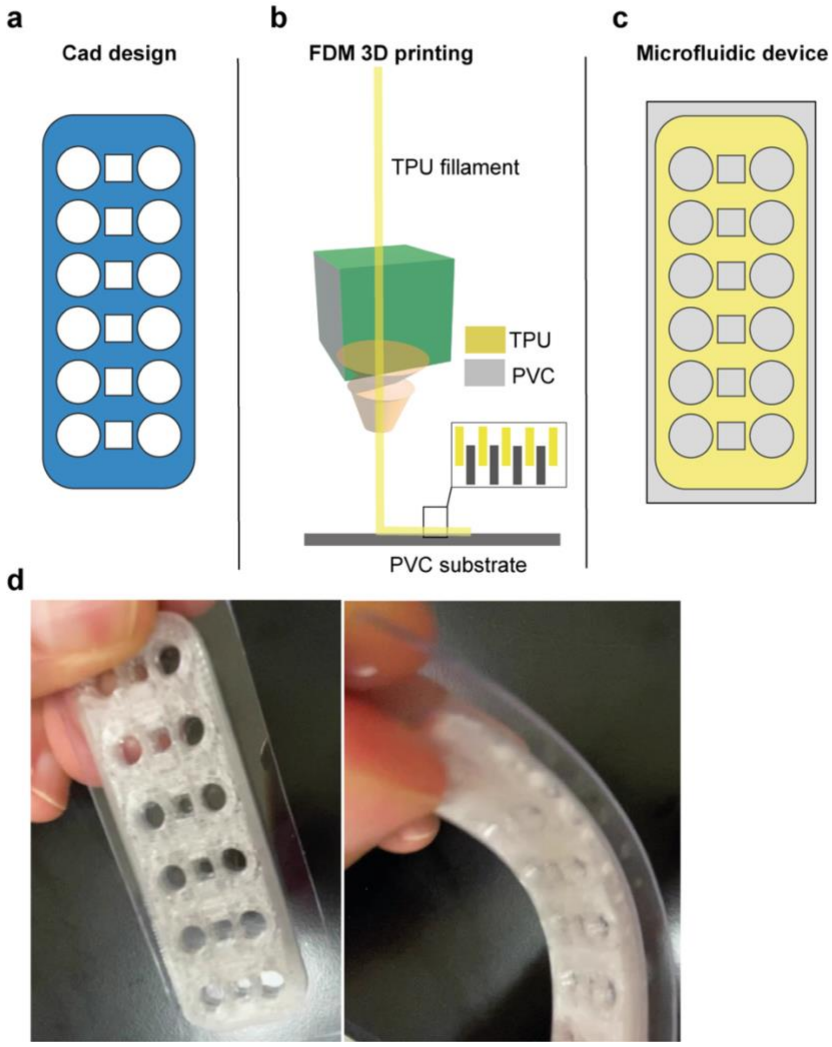


**Supplementary Figure 1. Design and fabrication of the 3D-printed microfluidic device.** (a) CAD layout showing the device design with central circular organoid chambers and side reservoirs. (b) Schematic of fused deposition modeling (FDM) 3D printing process using thermoplastic polyurethane (TPU) filament deposited directly onto a transparent polyvinyl chloride (PVC) substrate. Inset shows layered filament deposition. (c) Final schematic of the assembled TPU–PVC microfluidic device with six parallel culture units. (d) Photographs of the fabricated device showing optical transparency and flexibility of the TPU–PVC structure.
